# Early *NOTCH1* mutation is positively selected but epistatically suppresses evolution of later esophageal squamous-cell carcinoma drivers

**DOI:** 10.1101/2023.11.03.565535

**Authors:** Kira A. Glasmacher, Jeffrey D. Mandell, Mia Jackson, Nic Fisk, Vincent L. Cannataro, Jeffrey P. Townsend

**Affiliations:** Department of Biology, Emmanuel College, Boston, MA; Program in Computational Biology and Bioinformatics, Yale University, New Haven, CT; Department of Cell and Molecular Biology, University of Rhode Island, Kingston, RI; Department of Biostatistics, Yale School of Public Health, New Haven, CT; Department of Ecology and Evolutionary Biology, Yale University, New Haven, CT; Yale Cancer Center, Yale University, New Haven, CT

**Keywords:** Somatic evolution, Esophageal squamous-cell carcinoma, Selection, Epistasis, *NOTCH1* mutation

## Abstract

**Background:** Somatic mutations commonly accumulate in histologically normal tissues and contribute to cancer development. However, many mutations that are common in normal tissue are also found in cancers from the same organ, making their role in malignant progression unclear. To address this, we asked whether such mutations increase cell proliferation and survival to different degrees at different steps of esophageal tumor development, and whether they alter selection for other driver mutations.

**Methods:** We used a quantitative evolutionary framework to distinguish underlying mutation rate from selection across two steps of esophageal development: from organogenesis to clonal histologically normal esophageal epithelium, and from that tissue to esophageal squamous-cell carcinoma. Analyzing sequence data from 2171 samples, we estimated step-specific selection on recurrent somatic mutations, corresponding to increased cellular division and survival, and tested whether mutations in one gene earlier in the trajectory changed selection for mutations on another. We additionally examined somatic copy-number alterations and single-cell transcriptomic profiles to contextualize these evolutionary patterns within broader genomic and cellular changes during progression.

**Results:** *NOTCH1* mutations were strongly selected during the clonal expansion of histologically normal esophageal epithelium, explaining their high prevalence in that tissue. However, for the first time, we show that there is little to no positive selection for *NOTCH1, NOTCH2*, and *FAT1* mutations during progression from clonal histologically normal esophageal tissue to esophageal squamous-cell carcinoma in humans, leading to a conclusion that these alterations promote clonal expansion in normal tissue, but do not drive malignant progression from established normal clones. Moreover, we provide a somatic genetic basis for this step-specific effect: we demonstrate for the first time that mutations in *NOTCH1* exhibit antagonistic epistasis with mutations of *TP53* and *RB1*, reducing selection for these key tumor suppressor alterations during tumor development. Consistent with this model, copy-number alterations associated with later ESCC progression were more strongly selected in *TP53*-mutant samples, supporting the idea that *TP53* disruption promotes a genomic context more permissive for malignant evolution.

**Conclusions:** Early somatic mutations can promote clonal expansion in normal tissue without promoting cancer, and in some cases may limit progression by reducing selection for later driver events. *NOTCH1* and other genes can shape evolutionary trajectories in ways that ultimately constrain malignant progression. By separating mutation rate from selection, quantifying step-specific genetic interactions, and considering broader changes in genomic and cellular context, our study shows that the effects of recurrent mutations depend strongly on disease stage and mutational context—what promotes clonal expansion in normal tissue may later impede growth or survival in tumors. These insights underscore the need for precision strategies that account for the shifting fitness landscape across premalignant and malignant stages, informing early detection, prevention, and therapeutic prioritization.

## Background

Somatic mutations accumulate as cells divide and are exposed to endogenous and exogenous mutagens. Many of these mutations have little to no effect on cell fitness, whereas others increase cell proliferation and survival and thereby contribute to benign or malignant growth [1]. The role of somatic mutations in cancer have been studied at the level of whole genomes [2], individual genes [3], and even specific nucleotides or amino acid substitutions [4]. Sequencing studies of histologically normal and precancerous tissues have further shown that many mutations common in cancers are also frequent in their presumed precursors—for example, in atypical hyperplasia preceding uterine endometrioid carcinoma [5]. However, mutation prevalence alone is insufficient to determine whether a variant promotes progression to cancer, because prevalence reflects both how often a mutation appears in cells and how strongly it contributes to cell proliferation and survival once it occurs. Accurate inference of selection therefore requires disentangling mutational and selective processes that underlie evolutionary dynamics across successive stages of disease development. In this setting, scaled selection coefficients, a measure of how much mutations increase cell proliferation and survival, should be estimated in a step-specific manner, using normal and malignant tissues from the same organ, distinguishing mutations that contribute to early clonal expansions from those that drive later cancer progression.

Detection of somatic variants in histologically normal tissues has historically been challenging because variants are often present in normal tissue samples at low allele frequencies due to the heterogeneous stem-cell composition of normal tissues. Recent technical advances, including deep sequencing of small biopsies containing clonal populations [6–9] and sequencing of microscopically defined clonal units [10,11], have enabled the identification of substitutions that are present in only a small fraction of cells. To infer the contribution of driver mutations to tumorigenesis, some studies have relied on variant prevalence among samples [10,12]. However, prevalence alone conflates the distinct influences of mutation rate and strength of selection: gene substitutions may be prevalent in normal or tumor tissue not because they are functionally important for cell survival or proliferation, but simply because they mutate more frequently in individual progenitor cells. Varying prevalences of somatic substitutions can arise from quantitative differences in underlying mutation rates that are unrelated to biological function in oncogenesis. Statistical models for driver detection have striven to account for these underlying differences across the genome by martialing both gene-level synonymous mutation rates and gene-level covariates [13]. Techniques using synonymous substitutions can quantify an underlying “neutral” mutation rate, examine their frequency relative to nonsynonymous substitutions to detect drivers [13] and quantify gene-level selection [3,8]. Indeed, using such methods, recurrent mutations in cancer-associated genes have been identified in histologically normal skin, esophageal, colorectal, and endometrial epithelia [6,9,10,14], as well as in clonal hematopoiesis [15,16].

Application of methods that detect somatic selection to skin epithelia has revealed strong positive selection for mutations in cancer-associated genes such as *NOTCH1* and *TP53* [17]. However, the presence of a positively selected mutation in normal tissues does not necessarily imply that these mutations contribute to the development of cancer. Some mutations may expand in otherwise normal cell populations due to selective advantages unrelated to malignancy, then “hitchhike” on subsequent, independent tumorigenic mutations to become somewhat prevalent in cancerous tissues. Moreover, the functional impact of a mutation can change over the course of tumor evolution as other mutations accumulate. This phenomenon, known as epistasis, describes a situation in which the selective effect of one mutation depends on the presence or absence of another. Such interactions are synergistic when one mutation increases selection for a subsequent driver event, and antagonistic when one mutation decreases selection of another. These mutation-by-mutation interactions can therefore alter which variants drive different stages of disease development [18]. The biological and clinical relevance of epistatic interactions is supported by large-scale functional genomic studies, including the Cancer Dependency Map, which show that gene dependencies and therapeutic vulnerabilities vary markedly across genetic backgrounds [19,20]. Recognition and understanding of this context-dependence is essential to elucidation of the roles of somatic mutations in tumorigenesis, and has crucial implications for informing therapeutic targeting, drug development, and clinical translation [21–24].

Why some mutant cell populations remain benign while others progress to cancer is still not well understood. A common expectation is that mutations in driver genes should become more frequent as tissues progress from benign to premalignant to malignant states [25,26]. However, several observations challenge this view. For example, BRAF V600E substitutions have been reported in 70–88% of benign nevi but only 40–45% of melanomas, and HER2 expression is more common in ductal carcinoma in situ than in invasive breast cancer [27]. These “inverted” patterns suggest that some mutations may support clonal expansion in non-malignant tissue without promoting progression to invasive disease—and in some cases, may even limit it.

The Notch signaling pathway is an evolutionarily conserved cell-signaling system mediated by four transmembrane receptors (NOTCH1–4) that interact with Delta-like or Jagged ligands. It regulates processes such as cell-fate decisions, differentiation, and is particularly important for preserving the structure and homeostasis of adult epithelia [28,29]. In cancer, however, its effects are strongly context-dependent: Notch signaling can promote tumor growth, acting as an oncogene in T-cell acute lymphoblastic leukemia and some breast cancers, but it can also act in a tumor-suppressive manner in myeloid malignancies and squamous-cell carcinomas of the skin, head, and neck [28,30].

Higa and DeGregori [31] proposed that *NOTCH1* mutant cells in the esophagus might have a lower likelihood of accumulating mutations that drive malignant progression because they have reached distinct adaptive fitness peaks associated with clonal expansions of the normal esophageal epithelium. Specifically, a reduction or loss of NOTCH1 function may provide a growth advantage in normal esophageal epithelium, where nonsense and essential splice mutations have exhibited especially strong positive selection [32,33]. However, the advantage conferred by NOTCH1 loss in normal epithelium may not persist once cells begin progressing toward malignancy, potentially reducing the relative success of *NOTCH1*-mutant clones in tumors [34]. Indeed, a recent mouse study showed that NOTCH1 mutations can drive clonal expansion in normal mouse esophageal tissue and impair mouse tumor formation [35]. Quantification of selection in human normal esophageal tissue and esophageal tumors, while accounting for differences in underlying mutation rate, could therefore clarify the role of *NOTCH1* and related genes in early clonal expansion and later cancer development in the human esophagus.

Our analytical framework estimates selection by comparing observed nonsynonymous mutation counts to neutral expectations derived from synonymous mutations and local sequence context. This approach distinguishes mutations that increase cell proliferation and survival from those that are simply frequent because they appear often in cells due to their genomic context. We extend this framework across sequential stages of disease development by estimating mutation rate and selection separately in distinct evolutionary steps, enabling determination not only of whether a mutation is selected, but also at what point during tumorigenesis its selective effects may change.

Here we investigate the selective effects of somatic single-nucleotide mutations in clonal histologically normal human esophageal (CHNE) tissue (generally considered to be “normal” tissue [7,8,36]) and esophageal squamous-cell carcinoma (ESCC). We derive a novel step-specific analytical framework to estimate both underlying mutation rates and scaled selection coefficients for mutations in ESCC-associated genes in CHNE and malignant tissues. To assess how mutations in one gene may alter selection for mutations in another, we also analyzed pairwise selective epistatic interactions among frequently mutated genes, including *NOTCH1* and *TP53*. To place these mutation-level evolutionary patterns in a broader developmental context, we performed complementary analyses of copy-number alterations and single-cell transcriptomic data. This integrative approach enabled us to determine how mutation rate, selection, and genetic interactions together shape clonal expansion in normal human esophagus as well as progression to human carcinoma.

## Methods

### Data acquisition and preprocessing

We assembled a dataset of 2757 esophageal samples, including 1016 CHNE tissues and 1741 ESCC tumors. ESCC data included 624 whole-genome and 879 whole-exome samples from a combined cohort spanning 17 datasets, including TCGA, ICGC, and other published studies [37]; together with 25 additional whole-exome ESCC samples from Liu et al. [38]; 10 whole-exome ESCC samples from Yuan et al. [36] 137 targeted-sequenced ESCC samples from cBioPortal [39–42], and 66 targeted-sequencing ESCC samples from Yokoyama et al. [7]. CHNE data included 34 whole-exome samples from Yuan et al. [36], 142 targeted-sequenced samples Yokoyama et al. [7] and 832 CHNE samples from Martincorena et al. [8]. In the Yuan et al. [36] cohort, we excluded eight samples originally labeled as “normal” because they showed severe dysplasia or carcinoma in situ and shared mutations with matched tumors.

For somatic variant calling in the CHNE cohorts, non-epithelial reference tissues were used as germline controls, including patient-matched blood [7,36] and normal muscle tissue [8]. To avoid redundancy from closely related clonal samples, we excluded biopsies in the Yokoyama and Martincorena datasets that appeared clonally related on the basis of shared mutations or close spatial proximity: adjacent biopsies were retained only if they shared no detected variants. To ensure robustness and improve generalizability, the dataset included samples from multiple populations (**Supplementary Table S1**), and studies processed through different sequencing and variant-calling pipelines (**Supplementary Table S2**), reducing dependence on any single methodology.

We identified ESCC-associated genes based on prior studies of somatic mutations in both CHNE epithelium and ESCC [43–49]. To ensure sufficient statistical power and broad relevance, we included only genes with mutations observed in at least 25 samples across our dataset. For selection analyses on single-nucleotide variants (SNVs), we excluded any mutations occurring within one or two base pairs of one another in the same tumor, as these variants may represent clustered events (multi-nucleotide substitutions) with different underlying processes compared to isolated SNVs. Reanalysis of the data without this exclusion did not alter our conclusions.

### Stepwise mutation rates

To estimate mutation rates across sequential steps of esophageal tumor development, we used a two-step framework spanning progression from esophageal organogenesis to CHNE tissue, and from CHNE tissue to ESCC. This framework enables inference of selection by comparing observed mutation counts to neutral expectations derived from synonymous mutations and sequence-context covariates. To estimate how mutation rates and selection shift over tumorigenesis, we applied this observed-versus-expected principle across two successive pathology-defined evolutionary intervals: Step 0→1 from esophageal organogenesis to CHNE tissue, and Step 1→2 from CHNE tissue to ESCC.

Because CHNE samples reflect earlier clonal expansions and ESCC samples reflect later malignant outgrowth, tissue type provides the temporal ordering used in this model. Individual biopsies may still harbor some degree of subclonality. However, we treated each sample as representing a dominant clonal expansion for purposes of selection inference. At typical whole-exome and whole-genome sequencing depths, variants present in only a single or very small number of cells are unlikely to be detected. Therefore, observed variants are expected to be present at moderate to high frequency within the sampled clone and were modeled as fixed.

To estimate gene-specific mutation rates separately for CHNE and ESCC samples, we used dNdScv [3] on all whole-exome and whole-genome samples, incorporating tissue-specific covariates [4]. We evaluated the fits of the negative-binomial model underlying dNdScv through its overdispersion parameter θ. In the ESCC cohort, θ = 5.23 indicated a suitable fit, with a variance below the mean expected mutation rate. In contrast, the CHNE cohort yielded θ < 1, suggesting overdispersion and greater-than-expected variance [3]. To assess the robustness of estimates obtained over subsets of the data, we performed a jackknife procedure by iteratively omitting individual samples and re-estimating model parameters. These jackknifed subsets yielded a satisfactory mean estimate of 1.86 for the overdispersion parameter. Moreover, analysis of all jackknifed subsets yielded results consistent with the conclusions of our study.

For each pathologically-defined tissue set, we used dNdScv to estimate the expected numbers of synonymous substitutions (*exp_syn_cv*) for each gene using its covariate-adjusted model, which provides an estimate of the neutral baseline mutation rate. Because synonymous substitutions are used here to represent the neutral background, these estimates provide gene-specific baseline mutation rates independent of selection. We then normalized these values by the number of samples in each cohort and by the number of synonymous sites per gene, yielding expected per-nucleotide mutation rates for gene *g, E*_0→1,*g*_ and *E*_0→2,*g*_. We assessed the proportion of mutations accumulated along the step from esophageal organogenesis to CHNE tissue (Step 0→1) for each gene, computing *p*_0→1,*g*_ = *E*_0→ 1,*g*_*/ E*_0→2,*g*_. Similarly, the proportion of mutations accumulated along the step from CHNE tissue to ESCC (Step 1→2) was assessed as p_1→2,*g*_ *=* 1 − *p*_0→1,*g*_. These step-specific proportions were used to model mutation and selection separately across the two discrete steps of somatic evolution and tumor progression.

We used cancereffectsizeR v. 2.10.2 [50] to estimate sample-specific mutation rates for designated sets of genomic sites and to quantify selection across stages of disease development. For each tumor *i* and each designated set of sites within gene *g*, cancereffectsizeR estimated the mutation rate *μ*_0→ 1,*g,i*_ across those sites. These estimates were informed by gene-level synonymous substitution rates (*E*_0→1,*g*_)—which serve as an indicator of the neutral background mutation rate—as well as by sample-specific mutational spectra derived from mutational signature analysis [4].

We then partitioned these mutation rates across two evolutionary steps: *μ*_0→ 1,*g,i*_ *= p*_*0*→ 1,*g*_ × *μ*_*0→* 2,*g,i*_, representing the mutation rate from esophageal organogenesis to CHNE tissue (Step 0→1), and *μ*_1→ 1,*g,i*_ *= p*_1→2,*g*_ × *μ*_0→2,*g,i*_, representing the rate during the progression from CHNE tissue to ESCC (Step 1→2). These step-specific mutation rate estimates were subsequently used to calculate selection coefficients separately for each step of tumor development.

To define the variants for gene-specific selection analyses, we identified recurrently mutated positions in each gene. For the oncogenes *PIK3CA* and *NFE2L2*, the analyzed set consisted of recurrently mutated sites. For the tumor-suppressor genes *FAT1, NOTCH1, NOTCH2, TP53, FBXW7*, and *RB1*, the analyzed set included both recurrently mutated sites and all nonsense/stop-loss variants [50]. Gene classification as oncogene or tumor suppressor followed Chakravarty et al. [51]. All reported mutation rates and scaled selection coefficients were calculated using these designated variant sets. To support transparency and reproducibility, metadata for all included somatic variants within the genes investigated are provided (**Supplementary Table S3**).

### Stepwise scaled selection calculations

In Mandell et al. [50], likelihood for the strength of selection *γ* was calculated for any single cohort of *N* tumors by maximizing

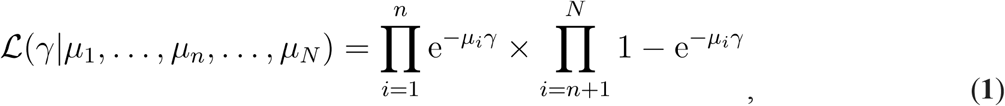

where μ_*i*_, 1 ≤ *i* ≤ *N*, is the mutation rate of a defined set of sites in sample *i*, and where *n* < *N* are defined such that any variant in the defined set of sites is absent in *n* samples and is present in samples *n* + 1 … *N*, and where the subscript *g* denoting gene identity is suppressed in **Eq. 1** and later equations for clarity of presentation.

To estimate step-specific scaled selection coefficients within a two-step sequential model of tumorigenesis, we elaborated the likelihood formulated by **Eq. 1** to differentiate Step 0→1 and Step 1→2:

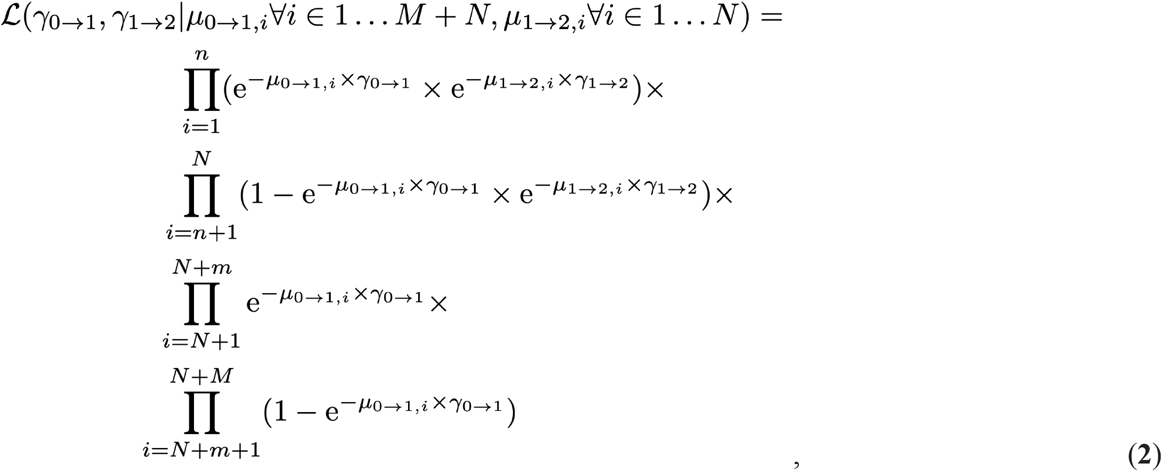

for a dataset with *M* CHNE samples, of which *M* − *m* exhibit a substitution and *m* do not, and where there are *N* ESCC samples in total, of which *N* − *n* exhibit a substitution and *n* do not. The integer variable *i* above is used to index CHNE samples *i* = 1 + *N*, …, *m + N, …, M + N* and ESCC samples *i* = 1, …, *n, …, N*. The first product term in **Eq. 2** represents ESCC samples where all variants in the designated set of variants are absent. The second product term represents ESCC samples where any variant in the designated set is present. The third product represents CHNE samples where all variants in the designated set of variants are absent. The fourth product represents CHNE samples with any of the variants in the designated set present.

The likelihood formulation in **Eq. 2** can be generalized to accommodate any number of sequential steps of tumorigenesis. For any number of steps *S*,

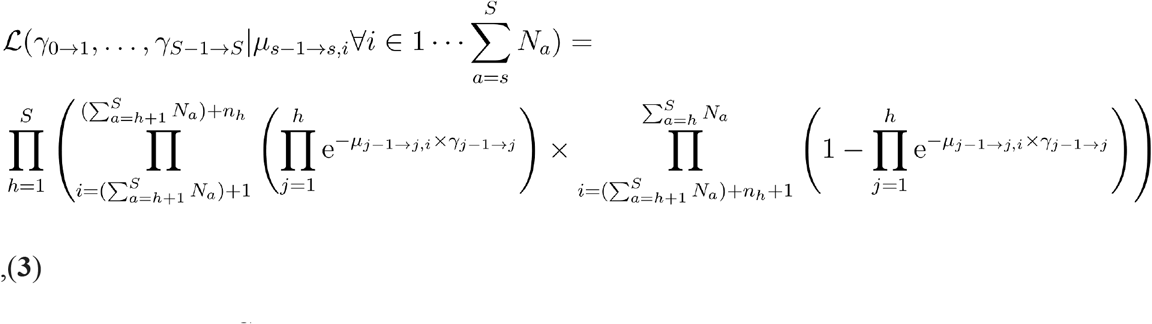

where *s* = 1, …, *S* denotes the step of progression toward cancer, and *S* corresponds to the last step present in the data. For each step *s*, there are *N*_*s*_ samples: *n*_*s*_ without a variant and *N*_*s*_ − *n*_*s*_ with a variant in the designated set of variants. The product index *h* specifies the step of a sample, *i* specifies the sample index, and *j* functions as an index that breaks down all *h* steps of a sample’s trajectory into the distinct stepwise processes. For each step, the samples are indexed by

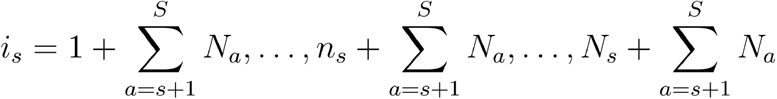

resulting in

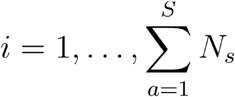

across all steps. By multiplying over all transitions in the trajectory, this likelihood captures the step-specific selection acting across a multi-step evolutionary process. This formulation enables flexible inference of selection coefficients in settings where data from intermediate steps—such as precancerous lesions—are available.

To determine whether scaled selection coefficients differed significantly across the steps of tumorigenesis, we performed a likelihood ratio test for each gene. Specifically, we compared the fit of the stepwise model—which estimates separate selection coefficients for each step—to a nested model that assumes a single, constant selection coefficient across all steps. *P* values were calculated based on a *X*^2^ distribution for the likelihood ratio test statistic with one degree of freedom (*df* = *s* − 1, where *s* represents the total number of steps in the stepwise model) under the null hypothesis of uniform selection. For reporting purposes, all tests yielding *P* < 0.001 were denoted as such, without additional decimal precision.

### Mutation rates for epistatic selection

To maximize statistical power and to capture interactions across the full span of disease development, we performed a separate analysis using all sequenced samples—both CHNE and ESCC. Using the standard cancereffectsizeR workflow [50], we estimated sample-specific mutation rates at individual sites without dividing the developmental process into discrete steps. These sample- and site-specific mutation rates were then used to calculate expected probabilities of substitution for modeling pairwise genetic interactions between genes.

### Epistatic scaled selection calculation

To improve on mutual exclusivity or co-occurrence analyses commonly applied in cancer genomics [e.g. 52,53], we conducted a likelihood-based analysis of epistasis. This method accounts for commonalities of underlying mutation rates for some driver gene pairs that are driven by common underlying mutational processes [54]. We first assessed the likelihood of the data under a null model of no selective epistasis. In this model, the Poisson rate of substitution for mutation A in each tumor *i* was estimated as 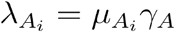 [54], where 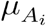 is the sample-specific neutral mutation rate, is the scaled selection coefficient for mutation A, and substitution is defined as acquisition of a mutation in a cell followed by its fixation in all or nearly all of the cells in a population. In the null no-epistasis model, the scaled selection coefficient *γA* was assumed to be constant, regardless of the presence or absence of a second variant B in another gene. Similarly, the selection coefficient for mutation B, *γB*, was inferred independently of the mutational status of A. In the absence of selective epistasis, substitutions are acquired independently with distinct selection coefficients *γA* and *γB*, which are therefore calculated separately as in **Eq. 1**.

For comparison, we analyzed the likelihood of the data under a model of pairwise epistatic selection. In this model, each tumor is characterized by four estimated rates of acquisition for each pair of variants: λ_*A*|∅_ and λ_*B*|∅_ for mutations A and B in the absence of the other substitution, and λ_*A*|*B*_ and λ_*B*|*A*_ for acquisition of A and B in the presence of the other substitution (for brevity, omitting the tumor index *i* arising from tumor-specific 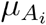.) These substitution rates were modeled as the product of sample-specific mutation rates and epistatic scaled selection coefficients. For this analysis, the underlying mutation rate is assumed to be constant for each gene across the full evolutionary trajectory, while selection on any variant may depend on the mutational background of another gene. We estimated selection by maximizing the likelihood function—multiplying over distinct probabilities of samples having only variant A, only variant B, neither variant, or both variants present—as in the pairwise epistasis approach of Alfaro-Murillo and Townsend [55]. We also used this model of pairwise epistatic selection to infer the order of mutation fixation through a continuous-time Markov chain that captures both possible directions of mutation fixation [55].

We used likelihood ratio tests to compare, for each gene pair, a model permitting epistasis versus a model assuming no-epistasis. Both models incorporated underlying gene-specific and trinucleotide-context mutation rates so that differences in co-mutation patterns would not be attributed to mutation rate alone. We then classified selective epistatic relationships as synergistic or antagonistic based on the statistical significance of the epistatic model fit parameterized by underlying gene and trinucleotide mutation rates and the directionality of the change in selection. A relationship was considered synergistic if the epistatic model predicted co-substitution patterns significantly better than the no-epistasis model, and predicted more co-occurrences than the no-epistasis model. Conversely, a relationship was classified as antagonistic when the epistatic model predicted co-substitution patterns significantly better than the no-epistasis model and predicted fewer co-occurrences than were expected in the no-epistasis model.

### Quantifying selection on copy number alterations

We quantified selection on somatic copy-number alterations (CNAs) in ESCC tumors from TCGA using a recently developed likelihood-based framework [56]. Allele-specific copy-number segments in ASCAT3 format [57] were processed and converted into copy-gain and copy-decrease events and mapped to disjoint gene-based genomic intervals on the hg38 reference genome. Large-scale anchored events were annotated using BISCUT-compatible definitions [58]. For each sample, background CNA rates were estimated from sample-specific CNA burden, distributions of copy segment lengths, and copy-number signature deconvolution [59,60]. Selection for each genomic interval was then estimated by maximum likelihood, comparing a model that permitted interval-specific deviation from neutrality with a model assuming no selection (**Supplementary Methods**).

For this analysis, we focused on a predefined panel of CNA loci previously implicated in ESCC progression (*FGF4, CDKN2B, SOX2, MYC, FOXA1, GBE1, CCND1, TEAD4, EIF4EBP1, EGFR, FGFR3, FZD6, PIK3CA, FAM135B, BCHE, SDHA* [61–63]). We first assessed co-occurrence between these CNA loci and the eight genes analyzed for SNVs. Because SNVs in *TP53* and *NFE2L2* were strongly associated with several CNAs, we performed selection analyses separately in strata defined by *TP53* and *NFE2L2* mutation status. For each candidate CNA locus, we report a locus-specific selection estimate obtained by comparing observed events with the neutral baseline rate. Most loci were evaluated individually without conditioning on selection at other CNAs. For chromosome arms containing multiple candidate drivers of ESCC (3q, 8q and 11q), we instead fit joint likelihood models including all candidates in the previously defined panel on that arm and report the conditional selection estimates from those models.

### Single-cell transcriptomic analyses

To provide an independent line of evidence for the tumor-intrinsic context of genes identified as under selection in our evolutionary framework, we analyzed publicly available single-cell RNA sequencing data from ESCC tumors and adjacent normal tissues (GSE160269 [64]). This dataset includes 208,659 single-cell transcriptomes from 60 ESCC tumors and four adjacent normal tissue samples generated using the 10x Genomics platform.

The CD45− and CD45+ UMI count matrices were merged into a single gene-by-cell expression matrix by taking the union of genes across both datasets and concatenating the corresponding cell barcode columns. Sample metadata were then assigned to individual cells using barcode identifiers and sample labels, enabling classification of cells as tumor or adjacent normal. Quality-control filtering retained cells with at least 200 detected genes, excluded genes detected in fewer than three cells, and removed cells with more than 20% mitochondrial transcripts. Expression values were normalized to 10,000 transcripts per cell and log-transformed. Highly variable genes were selected for dimensionality reduction, followed by principal component analysis, *k*-nearest neighbor graph construction, UMAP embedding and visualization, and Leiden clustering. Broad cell-type annotations were assigned using canonical marker genes for major cellular compartments, with each cell labeled according to its highest marker score.

To quantify variation in gene expression within each chromosome, genes were first ordered according to genomic position. The log-normalized expression values for each cell and gene were subtracted by the mean expression of that gene across all cells. For each cell, this expression profile was then smoothed along each chromosome by computing rolling means across the closest 100 neighboring genes. For each cell, we then quantified the standard deviation of the smoothed chromosome-level expression patterns to generate a chromosomal expression heterogeneity score. Differences between epithelial cell groups were assessed using Wilcoxon rank-sum tests.

## Results

### Cell-lineage expected mutational burdens differ among normal and tumor tissues

Across all whole-exome sequenced samples, the mean number of somatic substitutions was significantly higher in ESCC tumors than in CHNE tissues (*P* < 0.001). For every gene analyzed, the expected mutational burden per sample per synonymous site was lower along the evolutionary trajectory from esophageal organogenesis to CHNE tissue (*E*_0→ 1,*g*_) than in the corresponding evolutionary trajectory from esophageal organogenesis to tumor tissue (*E*_0→2,*g*_ ; **Fig. 1**). On average, the expected mutational burden in ESCC, estimated under a model of neutrality, was 38× greater than that in CHNE tissue. The distribution of these ratios was right-skewed, with several genes showing substantially larger differences. Among ESCC-associated genes, *NOTCH1* exhibited the largest contrast, with an expected mutational burden in ESCC that is 116 times greater than in CHNE tissue under neutral expectations.

**Figure 1.**
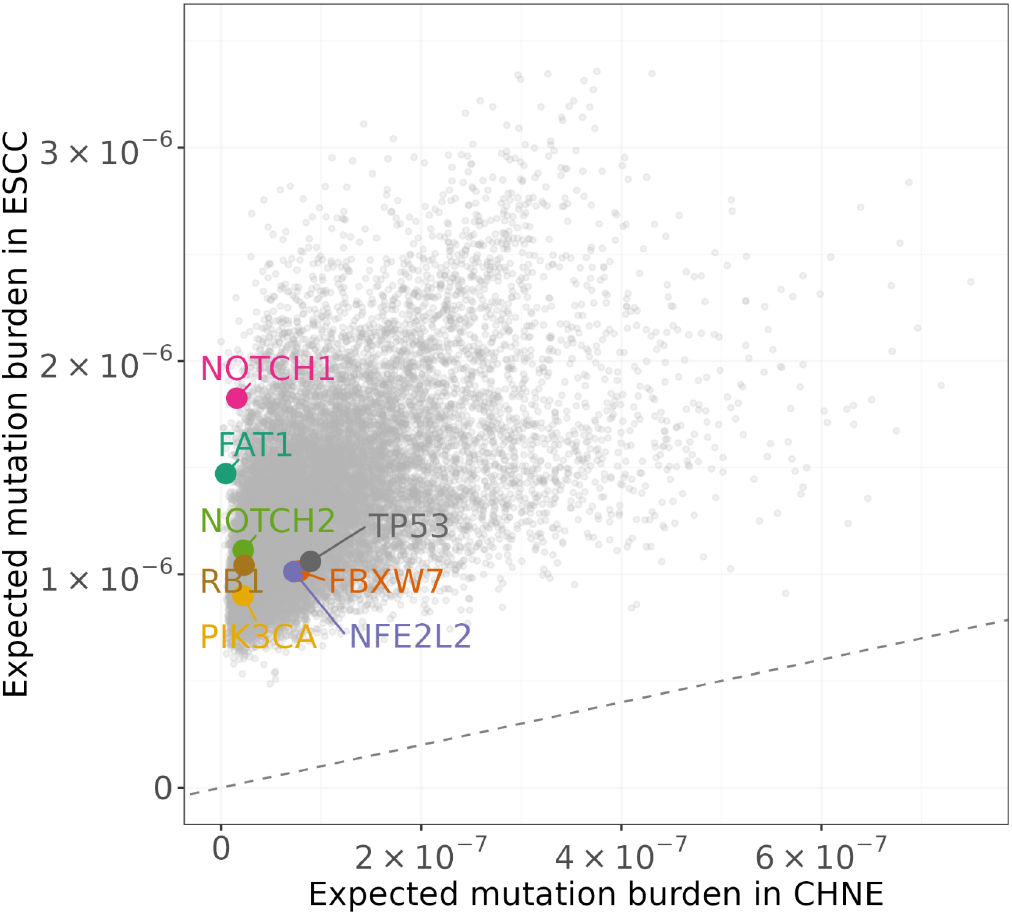
Expected mutation burden in genes in clonal histologically normal esophageal tissue and esophageal squamous-cell carcinoma. Gene-specific expected burden per sample per synonymous site were estimated from synonymous mutations and therefore reflect underlying mutation rate rather than selection. Each point represents one gene. Esophageal squamous-cell carcinoma-associated genes that are further examined in subsequent analyses are highlighted in color; all other genes are shown in gray. The dashed line indicates equal expected mutation burden in the two tissue types.

### Somatic mutations in NOTCH1, FAT1, and NOTCH2 are strongly selected during the transition to adult clonal histologically normal tissue, but are not selected during the transition to malignancy

*NOTCH1* mutations were detected in 61% of the sequences of CHNE tissue, compared to 23% of the sequences of tumor samples. One explanation for a difference in prevalence is a difference in underlying cellular mutation rate. However, the expected mutational burden per sample per site for *NOTCH1* under neutral assumptions was much lower in CHNE tissue (1.57 × 10^−8^) than in ESCC (1.83 × 10^−6^). Thus, the higher prevalence of *NOTCH1* substitutions in CHNE contrasts with the lower underlying expected mutational burden, a pattern consistent with a previous report [8]. Our analysis demonstrates, furthermore, that this discrepancy is the consequence of strong selection for *NOTCH1* driver substitutions along the step from organogenesis to CHNE. Moreover, there is either absent, negative, or very weak selection for *NOTCH1* driver substitutions along the step from CHNE tissue to ESCC (95% CI 0 ≤ γ ∼ 0 < 5.1; 0 < γ < 1 corresponds to selection against fixation; **Fig. 2**).

**Figure 2.**
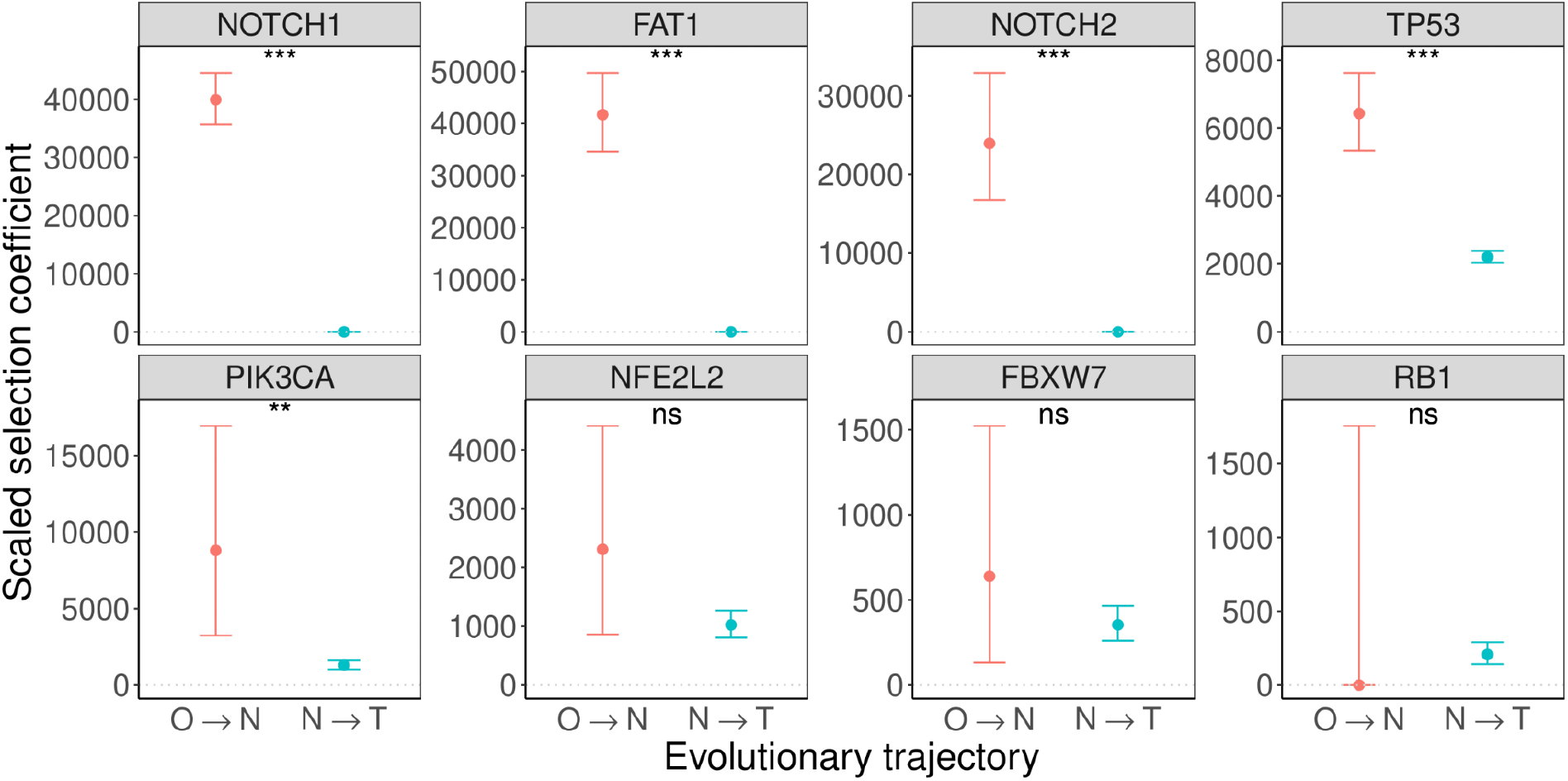
Step-specific selection on recurrently mutated genes associated with esophageal squamous-cell carcinoma in normal and malignant esophageal tissue. Scaled selection coefficients for mutations in genes previously reported as ESCC-drivers *NOTCH1, FAT1, NOTCH2, TP53, PIK3CA, NFE2L2, FBXW7*, and *RB1* are shown for two evolutionary steps: from esophageal organogenesis to adult clonal histologically normal esophageal tissue (O → N) and from adult clonal histologically normal tissue to primary esophageal squamous-cell carcinoma (N → T), with 95% confidence intervals. Genes are ordered by statistical significance for step-specific differences in selection, based on likelihood ratio tests comparing a model with separate selection coefficients for the two steps to a model with a single shared coefficient (***: *P* ≤ 0.001, **: *P* ≤ 0.01, *: *P* ≤ 0.05, ns: not significant).

*NOTCH1* is not the only gene in which observed mutation prevalence differed markedly from expected mutation burden. *FAT1* and *NOTCH2* showed similar patterns of low expected mutational burden and yet high prevalence in CHNE tissue, and conversely higher expected mutational burden but low prevalence in ESCC tumors. For *FAT1*, the expected mutational burden is 4.6 × 10^−9^ in CHNE and 1.5 × 10^−6^ in ESCC. In *NOTCH2*, the expected mutational burden is 2.2 × 10^−8^ in CHNE and 1.1 × 10^−6^ in ESCC. Despite these higher expected mutational burdens in tumors, *FAT1* and *NOTCH2* mutations were more common in CHNE than in ESCC (12.6% vs. 7.4% for *FAT1*; 7.2% vs. 2.0% for *NOTCH2*). Our analysis demonstrates that this discrepancy is the consequence of strong selection for driver substitutions in *FAT1* and *NOTCH2* during the transition from organogenesis to CHNE (34,608.1 < γ ∼ 41,718.9 < 49,743.2; 16,757.9 < γ ∼ 23,925.3 < 32,892.7), followed by absent, very weak, or negative selection for *FAT1* driver substitutions (0 ≤ γ ∼ 0 < 11.2) and *NOTCH2* driver substitutions (0 ≤ γ ∼ 0 < 38.2) during progression from CHNE tissue to ESCC.

Selection on mutated sites of *TP53, PIK3CA, NFE2L2*, and *FBXW7* was stronger in the transition from organogenesis to CHNE than in the transition from CHNE to ESCC (*P* < 0.001, *P* = 0.006, *P* = 0.200, and *P* = 0.496, respectively). Even so, these mutations were less prevalent in CHNE tissue compared to ESCC (*TP53*: 25.8% versus 73.0%; *PIK3CA*: 0.9% versus 6.2%; *NFE2L2*: 0.7% versus 6.4%; *FBXW7*: 0.7% versus 3.8%). This higher prevalence in ESCC can be attributed to continued substantial selection for new driver variants in these genes during progression from CHNE to ESCC. This constancy stands in contrast to *NOTCH1, NOTCH2*, and *FAT1*, in which driver-site selection was abrogated during progression from CHNE to ESCC.

### Selection on some driver mutations depends on the presence or absence of other driver mutations

Analysis of pairwise interactions across all samples, spanning the full course of disease development from organogenesis to tumor resection, revealed strong synergistic selective epistasis between *NOTCH1* and *NOTCH2* driver-site mutations (*P* < 0.001; **Fig. 3A**). In other words, the underlying mutation rates of *NOTCH1* and *NOTCH2* predicted fewer co-occurring cases under a model without interaction than were actually observed (5 expected vs. 14 observed). This excess co-occurrence was a consequence of stronger positive selection for *NOTCH2* mutations in the presence of *NOTCH1* mutation than in *NOTCH1*-wild-type tissue (γ ∼ 2713 versus γ ∼ 302).

**Figure 3.**
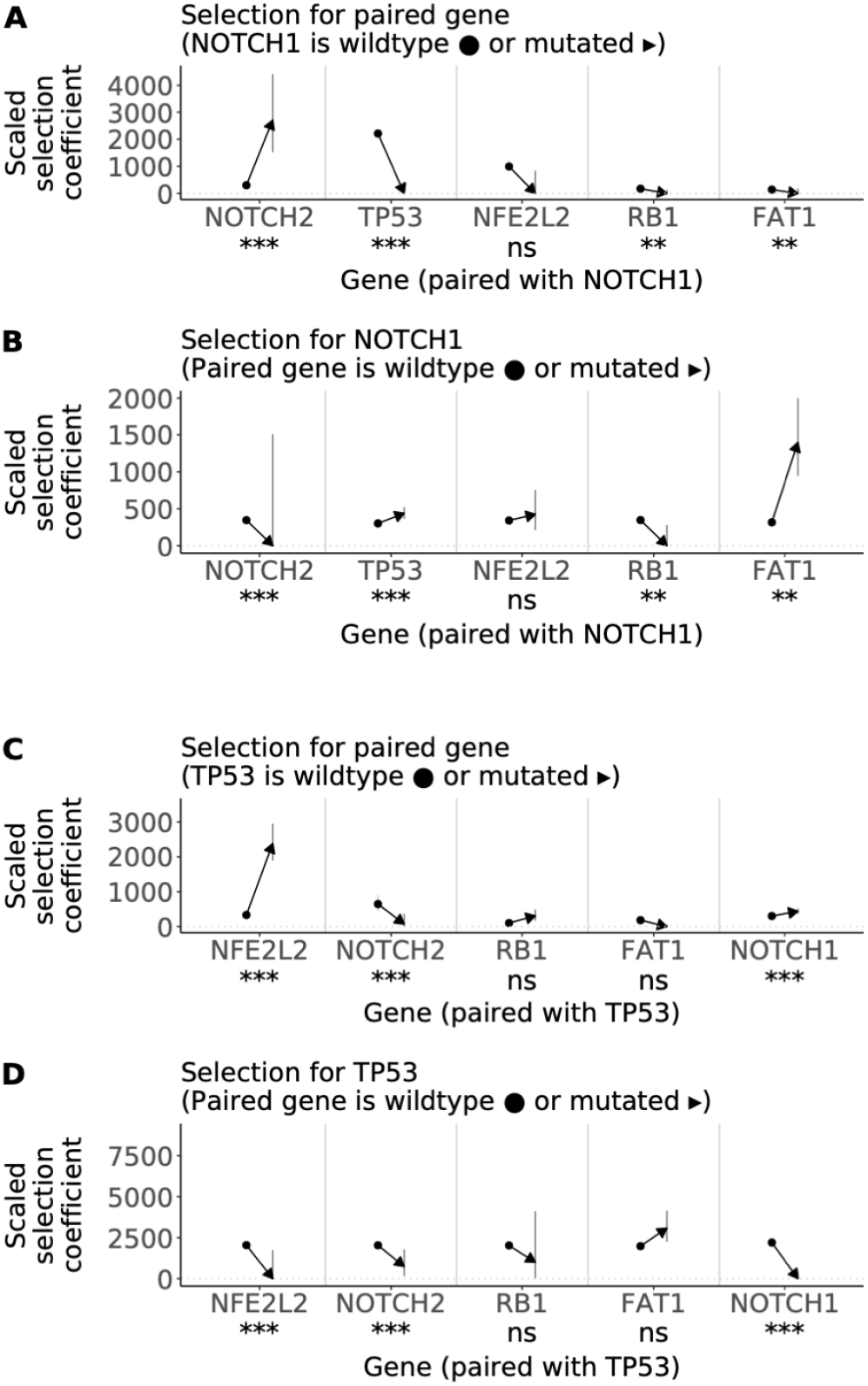
Pairwise epistatic effects on selection among recurrently mutated ESCC-associated genes. Context-specific scaled selection coefficients showing how the selective effect of a mutation changes according to the mutational status of another gene. These estimates account for underlying mutation rates, so differences between wild-type and mutant backgrounds reflect differences in selection rather than differences in mutation frequency alone. (**A**) Selection on mutations in *NOTCH2, TP53, NFE2L2, RB1*, and *FAT1* in *NOTCH1*-wild-type versus *NOTCH1*-mutant background. (**B**) Selection on mutations in *NOTCH1* in wild-type versus mutant backgrounds of *NOTCH2, TP53, NFE2L2, RB1*, and *FAT1*. (**C**) Selection on mutations in *NFE2L2, NOTCH2, RB1, FAT1*, and *NOTCH1* in *TP53*-wild-type versus *TP53*-mutant backgrounds. (**D**) Selection on mutations in *TP53* in wild-type versus mutant backgrounds of *NFE2L2, NOTCH2, RB1, FAT1*, and *NOTCH1*. Error bars depict 95% confidence intervals.

In contrast to the synergistic interactions, several gene pairs involving *NOTCH1* showed fewer co-occurring mutations than would be expected from their underlying mutation rates alone. This pattern was explained by antagonistic epistasis: in the presence of *NOTCH1* mutation, selection was reduced for driver mutations in *TP53* (*P* < 0.001), *RB1* (*P* < 0.01) and *FAT1* (*P* < 0.01; **Fig. 3A**). For *TP53*, a major driver of ESCC [7,8,65–68], selection decreased significantly from strongly positive in NOTCH1-wild-type tissue (95% CI 2080 < γ ∼ 2210 < 2345) to likely selection *against* fixation in *NOTCH1*-mutant tissue (0 ≤ γ ∼ 0 < 434; 0 < γ < 1 corresponds to selection against fixation). A similar pattern was observed for RB1, another established ESCC driver [67,68]: selection decreased from positive in *NOTCH1*-wild-type tissue (120 < ∼ 175 < 245) to likely selection against fixation in the presence of *NOTCH1* mutation (0 ≤ ∼ 0 < 139).

For *NOTCH1* and *RB1*, antagonistic epistasis was bidirectional: each mutation reduced selection for the other, regardless of which occurred first. In the presence of *RB1* mutation, selection on *NOTCH1* fell from positive in *RB1*-wild-type tissue (310 < γ ∼ 347 < 387) to likely selection *against* fixation (0 ≤ γ ∼ 0 < 282).

In contrast, interactions between *NOTCH1* and *NOTCH2, TP53*, or *FAT1* were asymmetric, with the direction of the effect depending on which mutation was already present. Selection on driver-site mutations of *NOTCH1* increased in the presence of a *TP53*-mutation (*P* < 0.001) or *FAT1* mutation (*P* < 0.01), but decreased in the presence of *NOTCH2* mutation (*P* < 0.001; **Fig. 3B**). Pre-existing NOTCH1 mutation reduced selection for *TP53* mutation—however, the reverse was not true: selection on driver-site mutations of *NOTCH1* was higher in *TP53*-mutant than *TP53*-wild-type tissue (364 < γ ∼ 439 < 524 versus 262 < γ ∼ 303 < 348). A similar pattern manifested for FAT1: NOTCH1 reduced selection for FAT1 mutation—however, selection on driver-site mutations of *NOTCH1* was higher in the presence of *FAT1* mutation (945 < γ ∼ 1399 < 1999 versus 281 < γ ∼ 317 < 356).

The relationship between *NOTCH1* and *NOTCH2* was more complex. Driver-site mutations of *NOTCH1* increased selection for driver-site *NOTCH2* mutations, whereas selection on *NOTCH1* mutations trended lower in the presence of *NOTCH2* mutations (0 ≤ γ∼ 0 < 1509 versus 309 < γ ∼ 346 < 386). The overall interaction between *NOTCH1* and *NOTCH2* was statistically significant (*P* < 0.001), a result that seems to be primarily driven by the substantial increase in selection of *NOTCH2* mutation in the presence of *NOTCH1* mutation (**Fig. 3A**), rather than by the reverse ordering (**Fig. 3B**).

Analysis of pairwise interactions across all samples also revealed strong synergistic epistasis between driver-site mutations of *NFE2L2* and *TP53* (*P* < 0.001; **Fig. 3C**). Specifically, selection on *NFE2L2* mutations was substantially stronger in the presence of *TP53* mutation than in *TP53*-wild-type tissue (1894 < γ ∼ 2382 < 2950; versus 224 < γ ∼ 333 < 472). As a result, co-occurrence of *NFE2L2* and *TP53* was more frequent than expected from underlying mutation rates alone. In contrast, a *TP53-*mutant background was associated with reduced selection for *NOTCH2* mutations (0 < γ ∼ 73 < 367; versus 444 < γ ∼ 644 < 897).

For *TP53* and *NOTCH2*, antagonistic epistasis was bidirectional: each mutation reduced selection for the other, regardless of which mutation occurred first. In particular, selection for *TP53* mutations was lower in the presence of *NOTCH2* mutation than in *NOTCH2*-wild-type tissue (*P* < 0.001; **Fig. 3D**). A different but likewise asymmetric pattern was seen for *TP53* and *NFE2L2. TP53* mutation increased selection for *NFE2L2* mutation, yet the reverse was not ture: in the presence of *NFE2L2* mutation, selection on *TP53* mutations fell from strongly positive in *NFE2L2*-wild-type tissue (1929 < γ ∼ 2048 < 2174) to likely selection against fixation in *NFE2L2*-mutant tissue (0 ≤ γ ∼ 0 < 1750).

### Selection on copy-number alterations

To assess how large-scale genomic alterations relate with SNV selective dynamics, we quantified selection on somatic copy-number alterations (CNAs) in ESCC. As an initial descriptive analysis, we evaluated co-occurrence between SNV mutation status and a curated set of recurrent ESCC-associated CNAs that have been previously reported (*FGF4, CDKN2B, SOX2, MYC, FOXA1, GBE1, CCND1, TEAD4, EIF4EBP1, EGFR, FGFR3, FZD6, PIK3CA, FAM135B, BCHE, SDHA* [61–63]). For most SNVs under selection in CHNE and ESCC, CNA co-occurrence patterns were weak or inconsistent. Two exceptions were apparent: tumors with *TP53* mutations exhibited increased frequency of co-occurrence (OR > 1) across multiple CNA loci than *TP53*-wildtype tumors, and tumors with *NFE2L2* mutations likewise showed elevated co-occurrence across several CNA regions.

To test whether these co-occurrence patterns reflected differences in CNA selection rather than differences in CNA frequency alone, we estimated scaled selection coefficients for each CNA locus separately within cohorts stratified by SNV mutation status. In analyses stratified by *TP53*, tumors with *TP53* mutation exhibited higher selection for many CNA loci of interest than *TP53*-wild-type tumors (**Fig. 4A**). In contrast, in analyses stratified by *NFE2L2*, selection for many CNA loci was similar or modestly lower in *NFE2L2*-mutant tumors than in *NFE2L2*-wild-type tumors (**Fig. 4B**). Finally, several CNA loci previously reported to be recurrently altered in ESCC [61–63] and speculated to be CNA drivers showed little evidence of positive selection; notably, despite recurrent CNAs and previous speculation about importance in ESCC due to significantly upregulated expression in amplified samples [62], *FGFR3* amplification exhibited no evidence of positive selection within any strata (**Fig. 4A–B**).

**Figure 4.**
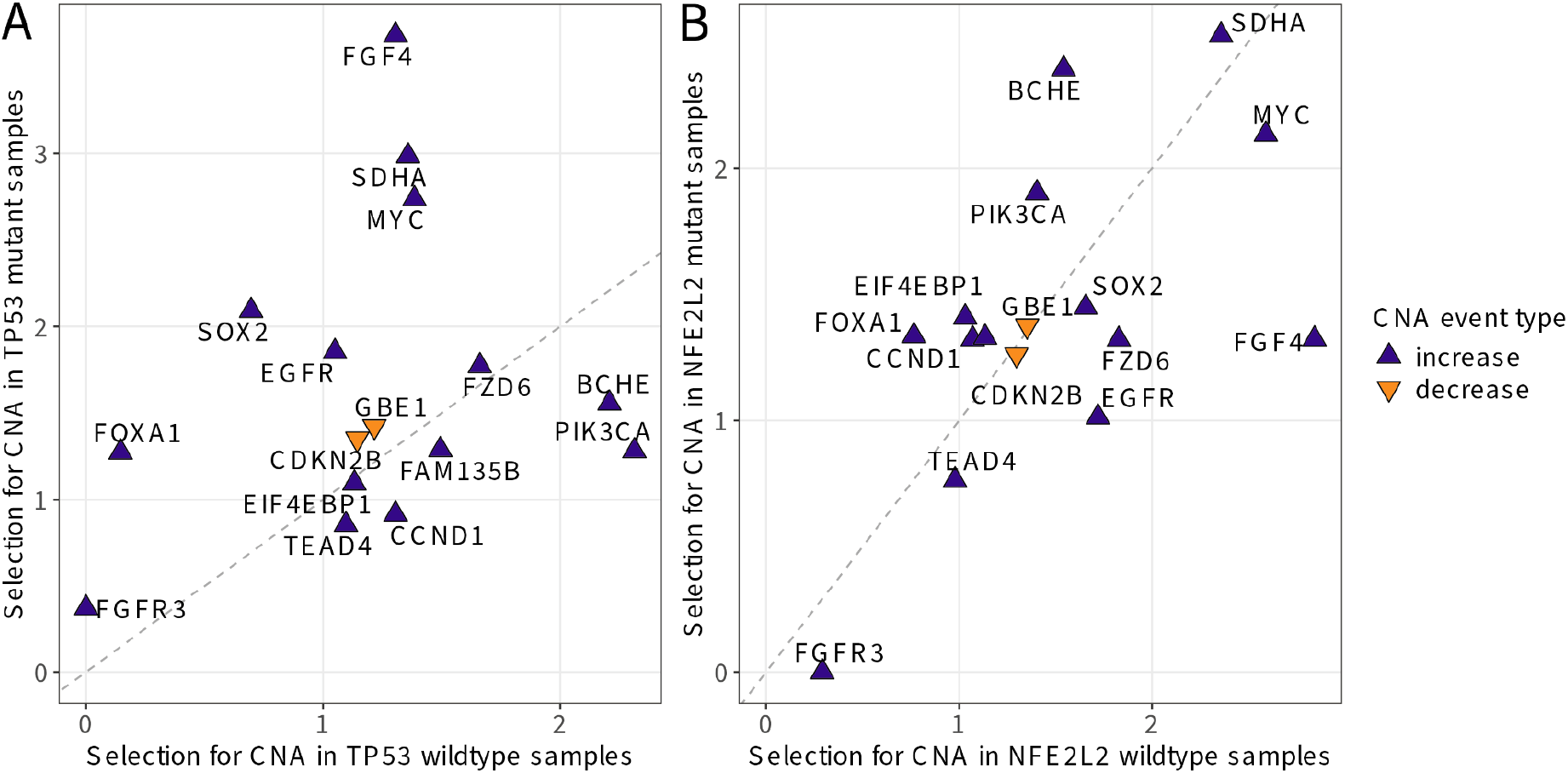
Selection on recurrent copy-number alterations in esophageal squamous-cell carcinoma, stratified by *TP53* and *NFE2L2* single-nucleotide mutant status. Each triangle represents one gene evaluated for CNA selection in wildtype tumors and in mutant tumors (blue upward-facing triangles: copy-number amplification; orange downward-facing triangles: copy-number decrease; dashed line: equal estimated selection in mutant and wildtype; neutral selection at 1) (**A**) *TP53*-mutant versus *TP53*-wildtype esophageal squamous-cell carcinomas. (**B**) *NFE2L2*-mutant versus *NFE2L2*-wildtype esophageal squamous-cell carcinomas.

### Single-cell transcriptomic analyses reveal increased genomic dysregulation during malignant progression and modest transcriptional shifts for genes under selection

To characterize the cellular expression of genes whose mutations were identified as under selection in our bulk DNA sequence analyses, we analyzed single-cell RNA sequencing data from 60 ESCC tumors and 4 adjacent normal tissues [64]. After quality-control filtering, 44,730 cells were retained for analysis. UMAP embedding and visualization revealed clear separation of epithelial, immune, and stromal populations, supporting robust cell-type annotation (**Fig. 5A**). Adjacent-normal samples were composed predominately of stromal, endothelial, and immune cells, whereas tumor samples contributed cells from multiple cellular populations, including a substantial epithelial population (**Fig. 5B**). This cellular resolution provides a framework for interpreting how genes identified as under selection in bulk tumor sequencing are expressed across tumor and microenvironmental cell types.

**Figure 5.**
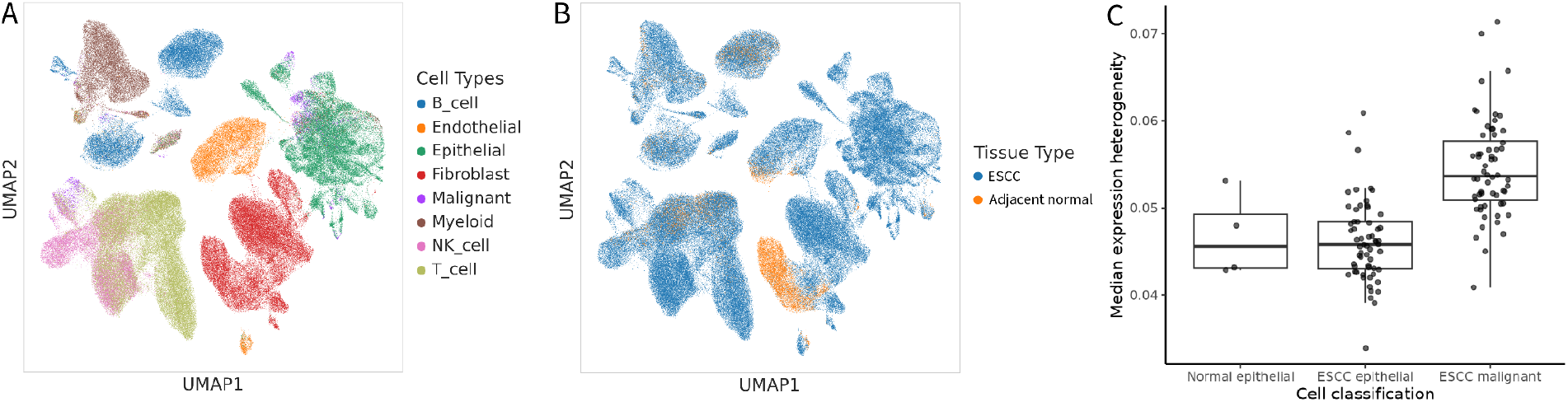
Single-cell transcriptomic context for genes identified in the bulk DNA sequence analyses. (**A**) UMAP embedding and projection of single cells colored by annotated cell type. (**B**) UMAP embedding and projection of single cells colored by tissue of origin. (**C**) Per-cell chromosomal median expression heterogeneity scores summarized by cell classification.

We quantified large-scale chromosomal expression heterogeneity at the single-cell level as a proxy for broad-scale transcriptional heterogeneity. Tumor epithelial cells exhibited significantly greater variability in gene expression along chromosomes than adjacent normal epithelial cells (Wilcoxon rank-sum test, *P* = 0.0079). Malignant epithelial cells exhibited even higher variability than other tumor epithelial cells (*P* < 2.2 × 10^−16^; **Fig. 5C**). Single-cell RNA sequencing does not directly measure mutation rates in the DNA. However, the observed increase in chromosomal expression heterogeneity and the proliferative signatures present in tumor epithelial cells are consistent with progressively increased genomic dysregulation during malignant progression, in line with elevated neutral background SNV mutation rates inferred in ESCC bulk sequencing data relative to CHNE, and a markedly higher prevalence of CNAs in ESCC compared with the rare copy-change events observed in CHNE.

For many individual genes whose mutations were identified as under selection in our bulk DNA sequence analyses, expression was higher in malignant ESCC cells than in other epithelial cell states. The magnitude of these differences was generally modest. The single-cell data therefore show transcriptional changes that are broadly consistent with the genes inferred to be under positive somatic selection inferred from bulk DNA sequencing. However, the results also show that mutations under strong selection do not necessarily produce large differences in transcript abundance. The modest expression differences may reflect context-dependent selective advantages or mechanisms that act independently of transcription.

## Discussion

Here we have developed a framework to quantify selection on somatic variants at successive stages of cancer evolution, moving beyond static models to capture the dynamic, stepwise forces shaping tumorigenesis. Applying this model and a complementary analysis of pairwise epistasis to somatic DNA sequence data obtained from 430 adult clonal histologically normal human esophageal tissue samples and 1,741 primary esophageal squamous-cell carcinomas, we estimated step-specific scaled selection coefficients for mutations in key cancer-associated genes. The results revealed that selective pressures shifted markedly between early clonal expansion and later malignant progression. These results resolve a long-standing paradox: why mutations in canonical cancer drivers such as *NOTCH1* and *FAT1* are often more common in premalignant human tissue than in human tumors [7,8]. Rather than manifestation of a contradiction, this pattern reflects fundamental, step-specific changes in selection—where early mutations may confer a fitness advantage in normal epithelium, yet become neutral or even deleterious as tumorigenesis proceeds. These findings validate the crucial importance of modeling evolutionary context and including premalignant states in research studies [69] when interpreting the somatic mutation patterns in human cancer [70,71].

Martincorena et al. [8] first reported strong positive selection on *NOTCH1* mutations in human clonal histologically normal tissue and speculated that such mutations might lose their selective advantage during malignant progression. Abby et al. [35] further investigated this idea in murine studies, suggesting that early *Notch1*-driven clonal expansions might suppress tumorigenesis in humans. Yet the somatic evolutionary dynamics underlying this paradox—where a putative driver mutation becomes less prevalent during malignant progression—have remained unresolved [27]. Our analysis provides direct evidence from human tissues that clarifies this discrepancy. We show that *NOTCH1* mutations are under strong positive selection during the transition from esophageal organogenesis to clonal histologically normal esophageal tissue, driving early clonal expansion, but are subsequently selected against during progression to ESCC and may instead be selected against.

This shift in somatic selection is especially striking because it occurs despite an increasing expected mutation rate as CHNE evolves into ESCC. Specifically, *NOTCH1* is expected to mutate more often in individual cells during progression to ESCC than during the earlier transition from organogenesis to CHNE. Yet *NOTCH1* mutations are more prevalent in CHNE than in ESCC. Accordingly, our analysis demonstrates for the first time in humans that the higher prevalence of driver-site substitutions in *NOTCH1* in CHNE samples must be a consequence of strong positive selection in clonal expansion whereas during progression from CHNE to ESCC, new *NOTCH1* mutations are no longer positively selected and may instead be selected against. Thus, despite a higher neutral expected mutational burden in ESCC, *NOTCH1* mutations become less common rather than more common. This striking reversal in selection across sequential steps during oncogenesis explains prior observations and reframes *NOTCH1* mutation not as a persistent driver of cancer, but as a context-dependent event that is advantageous in normal tissue, yet disadvantageous in the fitness landscape of tumorigenesis that can follow.

We identified strikingly similar patterns of mutation and selection in erstwhile ESCC driver genes *FAT1* and *NOTCH2 [39,72]*. Mutations in both genes are strongly positively selected during the transition from organogenesis to CHNE, yet newly arising mutations were not positively selected during progression to ESCC and may instead have been selected against—a result that has not previously been reported. Together with the findings for *NOTCH1*, these results demonstrate that mutations long considered canonical drivers of cancer may confer a selective advantage only in early, premalignant contexts. More broadly, they challenge the oft-made but poorly justified assumption that mutation prevalence directly reflects selective advantage. They reveal that the interplay between mutation and selection is profoundly shaped by developmental timing and somatic genetic context.

Indeed, *NOTCH1* and *TP53* mutations are both common along the developmental trajectory from organogenesis through CHNE to ESCC—driven by moderate to high mutation rates and strong positive selection—yet their co-occurrence in the same tumor is strikingly rare. In the absence of mutational correlations or epistatic effects, one would expect their frequencies to combine proportionally. Their mutual exclusivity suggests a biologically meaningful interaction effect. Experimental studies have shown that interactions between NOTCH signaling and p53 are highly dynamic and depend on cellular context, varying with cell state and tissue environment [73,74]. Our analyses now provide direct evidence from human tumors that this interaction has selective consequences: the presence of a *NOTCH1* mutation reduces selection for subsequent *TP53* mutation. Because *TP53* is among the most strongly selected drivers of progression to ESCC in *NOTCH1*-wild-type tissue, this antagonism may even help to limit malignant progression. We also report similar effects of *NOTCH1* mutation on driver-site mutations in genes such as *NFE2L2, RB1*, and *FAT1*, with statistically significant antagonistic epistasis for *RB1* and *FAT1* (*P* < 0.01). These findings suggest that *NOTCH1* mutation can promote early clonal expansion while simultaneously reducing selection for later driver events. Nevertheless, once *TP53* mutation has already occurred and altered the cellular state, subsequent mutations of *NOTCH1* may again become advantageous, obtaining new potential to contribute to cell proliferation and survival. More broadly, this dependence of selection on cellular state underscores the importance of cellular context in interpreting effects of somatic mutations.

In addition to antagonistic interactions, we have also identified synergistic epistasis, most notably between mutations of *TP53* and *NFE2L2*. Co-occurrence of these mutations has previously been reported in ESCC [75] and non-small-cell lung cancer [76]. Our analysis goes further by demonstrating that *TP53* mutation increases selection for subsequent *NFE2L2* mutations during progression to ESCC. This finding aligns with experimental evidence from breast cancer cell lines, where mutant p53 alters expression of NFE2L2 target genes and affects cancer cell survival [77]. This concordance between our inferred epistatic directionality and experimentally identified molecular mechanisms supports the validity of our data-driven evolutionary modeling. It also highlights its ability to capture and characterize functionally relevant gene-gene interactions in tumor evolution, and transforms a previously descriptive observation into a fundamental evolutionary insight: *TP53* and *NFE2L2* are not merely co-mutated due to common underlying mutagenic processes, but rather work together biologically to drive tumorigenesis in esophageal squamous epithelium.

The selective epistasis between *TP53* and *NFE2L2*, like many of the epistatic interactions we identified, is strikingly asymmetric: the selective advantage of one mutation depends on the presence of another, but the reverse effect is not the same. In this case, *TP53* mutation increases selection for subsequent *NFE2L2* mutations, whereas *NFE2L2* mutation reduces selection for subsequent *TP53* mutations. Such asymmetry may seem counterintuitive from a purely genetic perspective. However, this kind of asymmetry of epistasis has been both experimentally demonstrated [78] and conceptually grounded in evolutionary theory [79–82]. Mutations in key genes that are drivers of proliferation at different stages of disease such as in *NOTCH1* or *TP53* will alter the cellular state and, potentially, the surrounding tissue microenvironment [83–89]. These changes impose functional consequences, creating new positive and negative selective pressures that affect the fixation probability of subsequent mutations. More broadly, our findings confirm that somatic evolution is not a linear combination of effects, but a context-dependent process shaped by the sequence and interaction of mutations. Accurate reconstruction of tumorigenesis requires not just identifying which mutations arise, but also understanding the conditionality of selection that governs when and how they are favored.

Our epistatic analyses focused on interactions among point mutations. However, tumor evolution in ESCC is also shaped by large-scale genomic alterations. Our analysis of how the selective dynamics identified in SNVs intersect with patterns of selection on CNAs revealed CNA-SNV relationships that were gene-specific rather than uniform across driver SNVs. Most driver mutations exhibited limited association with CNA burden. However, *TP53* and *NFE2L2* were consistent exceptions. In particular, *TP53* mutation was associated not just with increased co-occurrence but also with higher inferred selection for CNAs recurrently affecting specific genes, supporting claims that *TP53* disruption is associated with increased accumulation of CNAs of different scales that are recurrent across ESCC tumors [61,62,90–93] and moreover that loss of *TP53* function acts as a permissive factor for large-scale genomic alterations in tumor progression [65,89].

Based on our SNV epistasis analysis, *NOTCH1* mutations are selectively favored in early clonal growth but exhibit antagonistic epistasis with *TP53*, reducing the selective advantage of *TP53*-mutant clones when *NOTCH1* is mutated. Because *TP53* mutation was also associated with increased selection for CNA drivers of cancer, one possible interpretation is that early *NOTCH1*-mutant clonal expansion may indirectly limit the emergence of *TP53*-driven CNA-enriched tumor lineages. We were not able to directly test whether *NOTCH1* mutation status alters CNA selection, because NOTCH1-mutant tumors were not frequent enough to provide sufficient power to support reliable stratified analysis. Future analyses will benefit from greater sample sizes.

Ideally, we would have been able to estimate step-specific CNA selection along the same evolutionary trajectory used for SNV analyses (CHNE → ESCC). However, currently available copy-number data for CHNE samples contained substantial gaps in genomic coverage that precluded robust inference of focal CNAs, which are critical for gene-specific CNA selection inference. Moreover, CHNE tissue harbored very few CNAs relative to ESCC [65], yielding limited power to estimate distinct CNA selection parameters from organogenesis to CHNE. These factors are consistent with the interpretation that most CNA events arise during malignant progression rather than during early benign clonal expansion, a finding that complements the results of our step-specific SNV analyses.

Co-occurrence patterns alone cannot distinguish whether enrichment of CNAs in particular SNV-defined subsets is driven by biological effects of CNAs versus incidental differences in underlying CNA event rates, stage composition, or other correlates of malignant progression. The observation from our stratified selection analysis that *TP53*-mutant tumors show broadly elevated selection for many CNAs is consistent with models in which TP53 disruption is associated with an evolutionary regime characterized by stronger fitness benefits of copy-number change and/or more efficient fixation of CNAs, aligning with established links of *TP53* with chromosomal instability [65]. In contrast, the absence of a similar elevation in *NFE2L2*-mutant tumors suggests that the elevated CNA co-occurrence observed for *NFE2L2* may reflect covariation with genomic instability or tumor context rather than systematically stronger selection on CNAs.

We found no evidence of positive selection for *FGFR3* copy-number increases despite previous reports suggesting *FGFR3* as a potential CNA-associated driver in ESCC [62]. This lack of selection highlights that some frequent CNAs may arise from chromosomal, subchromosomal, regional, and focal susceptibility to copy-number change rather than positive selection. In terms of its genomic context, *FGFR3* is near to a telomere, a position that can exhibit elevated background rates of copy-number change, potentially producing recurrent alterations with limited selective advantage.

Single-cell transcriptomic analysis of ESCC tumors and adjacent normal tissues showed that genes identified as under step-specific selection in the bulk DNA sequence variant analyses were robustly expressed within epithelial compartments. However, differences in transcript abundance were variable and did not track directly with estimated selection coefficients. Tumor epithelial cells also showed greater expression heterogeneity and stronger proliferative signatures than normal epithelium. These observations indicate that strong somatic selection on a gene does not necessarily correspond to large changes in steady-state transcript abundance, consistent with the possibility that many driver mutations act through signaling, protein function, or context-dependent regulatory effects rather than through large transcriptional shifts.

Interpreting selection on somatic mutations also requires consideration of the clonal microenvironment in which mutations arise. The normal and tumor samples analyzed here were treated as clonal expansions rather than as mixtures of many competing clones. In settings where multiple positively selected clones coexist, competition among them can reduce the likelihood that any one clone becomes dominant, a phenomenon known as clonal interference [94,95]. This competition lowers the probability of fixation for any competing clone, consequently reducing estimates of the strength of selection [96,97]. As a result, our own estimates of selection are likely to be conservative in absolute terms. However, this conservatism does not undermine our comparisons of relative strengths of selection across genes, because all genes within a given tissue context are expected to be subject to the same broadly competitive dynamics. Exceptions would require unusual biology, such as somatic variant clone cross-feeding or asymmetric interclonal toxicity. Accordingly, clonal competition can be expected to reduce observed mutation frequencies of substitutions and lower absolute estimates of the strength of selection; it should not compromise our ability to detect and quantify relative selective impact of mutations or to detect and quantify epistatic interactions between genes.

Individual mutations within the same gene can differ substantially in their biological effects and in the degree to which they are selected. Because our analyses were performed at the gene level rather than the single amino-acid variant level, each gene-specific estimate should be interpreted as an average of a spectrum of selective effects that occur on each variant within the same gene [1,4,98] Therefore to reduce dilution by likely neutral variants, we restricted inference to putatively selected variants based on recurrence and functional annotation, including recurrently mutated sites for oncogenes and recurrent and truncating variants for tumor suppressor genes. This restriction balances biological specificity with the statistical power required for step-specific estimation, since most individual variants are too rare for robust independent step-specific inference.

If there were sufficient inferential power to perform it, site-specific quantification would highlight specific sites within genes with modest and outsized roles in driving tumorigenesis. However, our assumption of a single cancer effect across genes is common in cancer evolution studies [3,8,99], and is especially reasonable for genes such as tumor suppressors where inactivation of function is achieved by a range of mutations with similar phenotypic consequences. Consistent with this expectation, highly recurrent variants in genes such as *TP53* showed site-level patterns among recurrent variants that qualitatively mirrored the gene-level selection trends, suggesting that aggregation did not obscure divergent variant-level behavior in these cases. As larger datasets become available, future studies should be able to estimate selection and pairwise epistatic selective effects at the level of individual variants, enabling a more fine-grained view of the architecture of mutational fitness landscapes in cancer evolution.

Patient population composition, genetic ancestry, germline and cultural effects are another potential source of bias in comparative analyses of somatic evolution based on specific tumor sequence datasets. Step-specific selection inference in our framework requires whole-exome or whole-genome sequencing to estimate baseline mutation rates. Among currently available CHNE datasets, sequencing of sufficient breadth is presently available primarily from a Chinese cohort, limiting fully ancestry-stratified step-specific inference. To assess robustness, we repeated key step-specific analyses within this cohort alone. Statistical significance was reduced due to smaller sample sizes. However, directional selection patterns across genes were qualitatively consistent. Likewise, across tumor cohorts drawn from multiple geographic regions represented in this study, we did not observe any outlier ancestry groups driving the reported gene-level selection trends. Larger datasets with more systematic ancestry representation will enable refined testing for differences in population-specific estimates of somatic selection and epistatic interactions.

At the gene level, we found suggestive evidence of antagonistic epistasis between *NOTCH1* mutants and driver-site mutations of *PIK3CA, NFE2L2*, and *FBXW7*, consistent with a broader pattern that *NOTCH1* mutations significantly attenuate selection on other, more common ESCC drivers. However, these interactions did not reach statistical significance at the current sample size (*P* > 0.05). Lack of statistical significance derived from limited sample sizes was especially evident for early, histologically normal tissues, where *NOTCH1* mutations are common but sequencing data remain sparse. As a result our ability to detect additional epistatic interactions in early steps of oncogenesis is limited. In particular, epistatic effects are likely to be context-dependent, varying across genetic backgrounds, microenvironmental conditions, and exposure histories: larger datasets, especially those performing deep sequencing of normal epithelium, and more stratified cohorts will likely reveal additional selective epistatic interactions that were not detectable in the present dataset [cf. 100,101]. Indeed, such datasets should also make it possible to move beyond pairwise interactions, empowering estimation of three-way, four-way, and higher-order combinations of mutations [55], revealing a more detailed view of how somatic mutations act together during cancer development via the fully nonlinear, multidimensional fitness landscapes navigated by evolving somatic cells.

These computational inferences of synergistic and antagonistic epistasis motivate experimental validation to establish their mechanistic basis. A range of complementary validation approaches could be pursued across model systems. For example, engineered cell lines carrying single and paired driver mutations could be used to compare effects on proliferation and clonal competition in controlled co-culture assays. Three-dimensional esophageal organoid models would enable these variant combinations to be tested in a more tissue-relevant epithelial setting, with lineage tracing and longitudinal sequencing to quantify clonal dynamics. *In vivo* studies in genetically engineered or xenograft mouse models could be used to investigate potential mechanisms of interactions affecting tumor initiation and progression.

At the same time, our results are derived directly from evolutionary patterns observed in large collections of human tumors and normal tissues, and therefore capture realized selection in the native human setting, including local tissue architecture, microenvironment, and immune context. As such, the model-based experimental systems and our population-level evolutionary inferences provide orthogonal and complementary lines of evidence: experimental systems can test specific mechanistic hypotheses under controlled conditions, whereas our approach measures the net selective consequences of mutation combinations as they occur in human disease.

Our analysis has revealed that selection on somatic mutations changes substantially between clonal histologically normal esophageal tissue and esophageal squamous-cell carcinoma, and that these changes are further shaped by gene-specific epistatic interactions. These insights into *NOTCH1, TP53, NOTCH2, FAT1*, and *NFE2L2*, coupled with their associated epistatic interactions, indicate that early clonal expansion and later malignant progression are governed by different selective conditions. They underscore the importance of modeling early cancer progression as a distinct evolutionary phase, governed by selective pressures that differ fundamentally from those in advanced tumors. A key clinical implication is that the mutations most prevalent in ESCC are not necessarily the most effective targets for early detection or prevention. By distinguishing underlying mutation rate from selection and quantifying how selection shifts across disease stages and in distinct somatic genetic backgrounds, our methodology enables a more stage-specific precision oncology strategy that is attuned to the timing and trajectory of tumor evolution. Earlier diagnosis and intervention can improve survival, preserve quality of life, and reduce treatment burden; accordingly, an understanding of when specific mutations promote progression—and why they instead constrain it in certain contexts—can guide development of more effective, precision approaches to risk management, prevention, and treatment.

## Conclusions

Our study provides new insight into how somatic mutations contribute to the evolutionary process underlying esophageal tumorigenesis. We introduce a novel step-specific modeling framework that explicitly accounts for variation in underlying mutation rates across the progression from esophageal organogenesis to clonal histologically normal esophageal tissue and subsequently to esophageal squamous-cell carcinoma. This approach enables precise inference of selective pressures at discrete steps of tumor development. Application of this model to sequencing data from normal and tumor tissues of the human esophagus, we showed that both mutation rate and selection vary substantially across this evolutionary trajectory. We further revealed, for the first time, that new mutations of *NOTCH1, NOTCH2*, and *FAT1* are strongly selected in CHNE tissue, but not during further progression to esophageal squamous-cell carcinoma, and that this shift is associated with antagonistic epistasis of *NOTCH1* with key tumor suppressors including *TP53* and *RB1*. These findings illuminate how somatic alterations can support clonal expansion in normal tissue while limiting later malignant progression, offering crucial insight for precision oncology which must take location along the oncogenetic trajectory and somatic genetic context into account when investigating, designing, and testing translational approaches for risk assessment, early detection, and stage-specific therapy.

## List of abbreviations

CHNE: Clonal histologically normal esophageal
ESCC: Esophageal squamous-cell carcinoma
SNV: Single-nucleotide variant

## Declarations

### Ethics approval and consent to participate

Not applicable.

### Consent for publication

Not applicable.

### Availability of data and materials

No new datasets were generated for this study. The datasets supporting the conclusions of this article are available through cBioPortal (https://www.cbioportal.org/study/; ID escc_ucla_2014), Synapse (https://www.synapse.org/; ID syn33401857), and the supplementary materials of Martincorena et al. [8], Yokoyama et al. [7], Yuan et al. [36], and Liu et al. [38].

All analyses were conducted using a fully reproducible computational pipeline that is freely available at https://github.com/Cannataro-Lab/ESCC_step_epistasis and on Zenodo (https://doi.org/10.5281/zenodo.15659607), which utilized cancereffectsizeR v.2.10.2 [50], dependent on R (version ≥ 3.5.0).

### Competing interests

The authors declare no competing interests.

### Funding

This research was supported by the Elihu Endowment at Yale to JPT and by NIH NLM 5T15LM007056-32 and NIH NCI 1F31CA257288-01 to JNF. KAG was also supported by the Emmanuel College School of Science and Health.

### Authors’ contributions

Conceptualization: VLC and JPT; Data curation: KAG and VLC; Formal Analysis: KAG, VLC, JPT; Funding acquisition: VLC, JPT; Investigation: KAG, MJ, NF, VLC; Methodology: KAG, JDM, NF, VLC, JPT; Project Administration: VLC, JPT; Resources: VLC, JPT; Software: KAG, JDM, NF, VLC; Supervision: VLC, JPT; Validation: KAG, VLC; Visualization: KAG, JDM, NF, VLC, JPT; Writing—original draft: KAG, MJ, VLC, JPT; Writing—review and editing: KAG, JDM, NF, VLC, JPT.

## Acknowledgements

We thank Christopher Cross for helping to organize early efforts to start this project, Stephen Gaffney for arranging early data acquisition, Jose Costa for discussion of the topic, and Wes Lewis and Elizabeth Perry for helpful comments on an early draft.

## Additional Files

“Supplementary_methods.pdf”

### Supplementary Methods

Description of preprocessing of ASCAT-derived allele-specific copy-number data, ploidy-aware classification of SCNAs, estimation of baseline copy-number event rates, and the likelihood-based framework used to infer selection on gene-level copy-number changes in ESCC tumors.

“Supplementary_table_1.tsv”

**Supplementary Table 1**. Source and classifications of all normal and tumor samples included in this analysis.

^*a*^ Study indicates the publication or dataset from which the sample was obtained for this analysis. This may differ from the original publication in which the sample was first described.

^*b*^ Normal_or_Tumor indicates whether the sample was derived from clonal histologically normal esophageal epithelium (CHNE) or tumor tissue (ESCC).

^*c*^ Coverage refers to the type of sequencing performed (i.e., targeted panel, whole-exome, or whole-genome).

^*d*^ Notes provide the original source study for samples drawn from the integrated cohort assembled by Li et al. (2022).

“Supplementary_table_2.xlsx”

**Supplementary Table 2**. Summary of sequencing targets and variant calling pipelines used across studies.

^*a*^ Study, publication year, and PMID refer to the original source for sequenced samples.

^*b*^ Samples indicate the number of normal and tumor samples processed in the cited study, as reported by the authors.

^*c*^ Average sequencing depth refers to the reported median or mean sequencing coverage per sample, when available.

^*d*^ Additional comments on methodology summarize study-specific quality filters, exclusion criteria, or other processing steps not captured in the preceding columns. “Supplementary_table_3.tsv”

**Supplementary Table 3**. Summary of called variants across all datasets within the eight genes of interest.

## Notes

https://github.com/Cannataro-Lab/ESCC_step_epistasis

https://doi.org/10.5281/zenodo.15659607

### Competing Interest Statement

The authors have declared no competing interest.

### Summary of Updates

New results: selection on copy number alterations and single-cell transcriptomic analyses; Revised figures throughout; Supplemental methods added

