## Supplementary Methods for "Early *NOTCH1* mutation is positively selected but epistatically suppresses evolution of later esophageal squamous-cell carcinoma drivers"

### Preprocessing, ploidy normalization and event classification

Somatic copy-number alterations (CNAs) were analyzed using allele-specific ASCAT calls, in which integer copy number is inferred from bulk tumor samples containing mixtures of normal and tumor cells. Starting from these ASCAT segment calls, we preprocess and classify copy-number changes, estimate sample-specific CNA burdens and baseline event rates, and then infer interval-specific selection from the observed distribution of copy-number changes across tumors in comparison to the neutral baseline rates (**Suppl. Methods Fig. 1**).

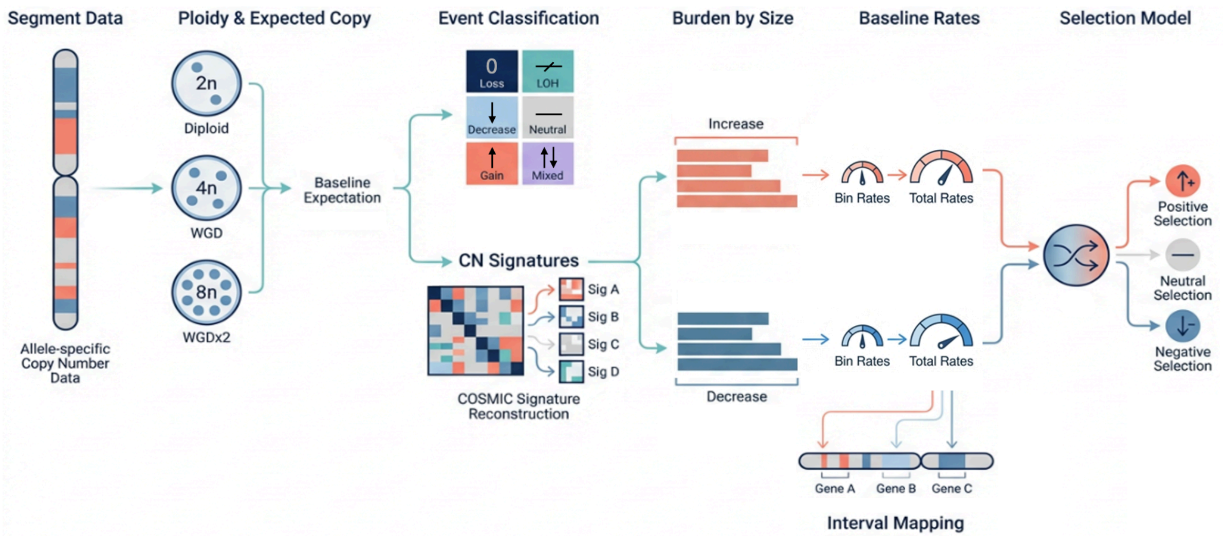


**Supplementary Methods Figure 1**. Schematic of CNA methods applied to ESCC tumors from the TCGA-ESCA project.

We analyzed allele-specific copy-number calls in ASCAT format [1] which reports segment-level major and minor copy numbers across the genome. ASCAT segments the genome into intervals defined by genomic coordinates and allele-specific minor and major copy numbers, with the minor copy number constrained to be less than or equal to the major copy number. For example, a diploid segment has major and minor copy numbers of 1 and 1. ASCAT output does not provide explicit chromosome phasing, so the reported major and minor copy numbers identify the larger and smaller allelic allelic copy states within a segment but do not specify which parental homolog carries each state.

ASCAT3 copy-number segments were obtained from the Genomic Data Commons for TCGA ESCC tumors, a subset of the TCGA-ESCA project, and analyzed on the hg38 reference genome. Segment coordinates were harmonized across samples by enforcing non-overlapping contiguous intervals within each chromosome; when the end coordinate of one segment coincided with the start coordinate of the next segment, we reduced the end coordinate of the upstream segment by one base pair to avoid overlap. For each chromosome, we defined the callable region as the intersection of genomic coordinates covered across all samples. Chromosomes without segment coverage over the p arm were annotated as lacking callable p-arm sequence.

We then estimated an even-integer ploidy state per sample (diploid, whole-genome duplicated (WGD), or twice-genome duplicated) using a length-weighted measure of total copy number adjusted for loss of heterozygosity (LOH), following Steele et al. [2]. For male samples, the X chromosome was excluded from this ploidy calculation. The inferred ploidy state was then used to define the expected major and minor copy numbers for each sample, enabling classification of the copy state of each segment relative to its expectation. Because the X chromosome is haploid outside pseudoautosomal (PAR) regions in male samples, the expected minor copy number was set to zero in non-PAR regions. If an ASCAT segment crossed a PAR boundary in a male sample, we split the segment at that boundary so that PAR and non-PAR portions could be evaluated against the appropriate expected copy-number states.

Segments were next annotated with copy change consequence according to how the major (nMajor) and minor (nMinor) copy numbers vary from expected major ($nMajor_{exp}$) and minor ($nMinor_{exp}$) copy numbers (**Suppl. Methods Table 1**).

**Supplementary Methods Table 1**. Copy-number consequences inferred from major and minor copy-number comparison to ploidy-adjusted expected major and minor copy numbers.

| nMajor | nMinor | Copy change consequence |
|---|---|---|
| 0 | 0 | Loss |
| > 0 | 0 | Loss of heterozygosity (LOH) |
| $0 < nMajor < nMajor_{exp}$ | $0 < nMinor \leq nMinor_{exp}$ | Decrease |
| $nMajor = nMajor_{exp}$ | $nMinor = nMinor_{exp}$ | Neutral |
| $nMajor > nMajor_{exp}$ | $nMinor \geq nMinor_{exp}$ | Gain |
| $nMajor > nMajor_{exp}$ | $0 < nMinor < nMinor_{exp}$ | Mixed |

After segment preprocessing, we called large-scale CNAs and classified them by copy-number consequence. We first identified whole-chromosome events and then arm-level events. To call a chromosome-level or arm-level alteration, we required segment coverage across at least 99% of the corresponding region. After these chromosome- and arm-level calls were assigned, we identified more localized large-scale events: telomere-anchored events, centromere-anchored events, and trans-centromeric events. Telomere-anchored events were defined as alterations beginning near a telomere, centromere-anchored events as alterations beginning near a centromere, and trans-centromeric events, as alterations spanning the centromeric region from the p arm into the q arm. These event classes were defined using breakpoint-based region definitions compatible with the BISCUT framework [3].

BISCUT was originally developed to identify genes showing evidence of positive or negative selection on CNAs by evaluating the distribution of anchored CNA endpoints [3]. In that framework, trans-centromeric CNAs are represented as pairs of centromere-anchored events. For the present analysis, we reannotated such "paired" events as single trans-centromeric events. Anchored events were enumerated explicitly, and their corresponding copy-number contributions were removed from overlapping segments before downstream modeling. BISCUT was used here as a source for a compatible event-definition scheme; it was not used here for *de novo* peak discovery.

**Genomic intervals for testing**

Our framework enables estimation of selection on copy-number changes across any predefined set of genomic intervals, such as fixed-width windows or cytobands. For this application, we used a gene-centered approach. We defined candidate intervals as non-overlapping gene-based genomic coordinates derived from reference genome hg38 transcript annotations, so that each genomic position was assigned to at most one tested interval.

For each chromosome, copy-number segments were mapped to these disjoint gene-based intervals. If a sample did not have complete coverage of a tested interval within the callable genomic region, that sample was excluded for that interval. When multiple segments from the same sample overlapped a given interval, we assigned the interval the copy-number state of the smallest overlapping segment, on the rationale that more focal events are likely to provide more gene-specific information for gene-level inference of selection.

**Model of CNA selection**

We formulated the CNA selection model to parallel, as closely as possible, the assumptions used in prior models of selection on single-nucleotide variant (SNV) effects [4,5]. For each interval and event type $t \in \{inc, dec\}$ (copy-number increase or decrease relative to expected ploidy), we observe a binary indicator $y_{s,i,t} \in \{0,1\}$, where $s$ indexes samples, $i$ indexes tested genomic intervals, and $y_{s,i,t} = 1$ indicates that at least one copy-change event of type $t$ affected interval $i$ in sample $s$. As in the SNV framework, we assume that event counts in each interval are Poisson-distributed with a constant rate $v_{s,i,t}$. However, the observed data are analyzed in binary form, distinguishing intervals with no event from intervals with one or more events. This binary representation reflects a key feature of bulk copy-number data: the final clonal profile observed in the ASCAT3 data may result from one or multiple occurrences of the same class of event during clonal expansion, but the segmented copy-number call records only whether that state is present in the dominant clone. We further assume that once a copy-number increase or decrease—relative to the sample ploidy—has occurred and become fixed in the clonal lineage, that copy-number state does not subsequently revert.

**Measuring CNA burden and calculating baseline event rates**

To quantify sample-specific burdens of copy-number increase and copy-number decrease events, we applied a second layer of CNA classification based on the feature set described by

Steele et al. [2]. This feature set, which has been adopted by COSMIC, was developed for copy-number signature analysis, with the aim of identifying reproducible copy-number signatures linked to underlying biological processes, analogous to mutational signature analysis for SNVs. An important distinction, however, is that SNVs can nearly always be expected to be the products of single mutational events, whereas observed copy segments are not events themselves but the cumulative result of one or more underlying copy-number changes.
Using this framework, all segments were classified according to segment size, total copy number, and heterozygosity, generating per-sample profiles of category counts. We then assigned COSMIC copy-segment classes to the pre-processed segment data so that copy-number signature extraction could be performed via MutationalPatterns [6].

To parameterize background CNA incidence, we estimated the propensity of each sample for copy-number increases and decreases separately within COSMIC-defined segment-length bins $b$ (0–100 kb, 100 kb–1 Mb, 1–10 Mb, >40 Mb). COSMIC specifies a distinct set of bins for homozygous deletions, but because these events were too rare to effectively model in our dataset, they were excluded from this burden calculation. Copy-number signature reconstruction across these classes produced per-sample weights, which we then used to quantify the burden of increases and decreases by size bin $B_{s,t,b}$.

Because COSMIC copy-number classes are defined by absolute total copy-number (for example, the 3–4 copy class) rather than by deviation from sample-specific expected ploidy, some classes are ambiguous as to whether they represent an increase versus decrease compared to ploidy in genome-doubled tumors. To account for this ambiguity, we partitioned these classes according to the empirical frequencies of the underlying integer copy states observed within each ploidy group.

When evaluating the likelihood of an observed copy-number call set, we included all called segments overlapping any tested interval. Because a single segment can overlap multiple intervals, its contribution to the likelihood may be modified by multiple selection terms. To specify the corresponding baseline rates in the likelihood model, we converted sample-level CNA burdens into event-level baseline rates.

Burdens ($B_{s,t,b}$) quantified how many copy-number changes of a given type and size range are present in a tumor sample. For numerical stability in maximizing likelihoods, zero burdens were replaced with a small positive floor derived from the lowest non-zero burden observed. To convert this burden into a genomic event rate, we adjusted for the typical size of events in that class, because longer CNAs are more likely to overlap any given interval. Specifically, for each sample, event type, and COSMIC length bin, we defined a representative event length $L_{s,t,b}$ as the median observed segment length among events of type $t$ and length bin $b$. If a sample had no events in a given bin, we used the cohort-wide median length for that bin. We also defined the callable genome length $G_s$ as the total length of segments with defined copy state in that sample. The bin-specific baseline event rate was then calculated as $R_{s,t,b} = B_{s,t,b}\frac{L_{s,t,b}}{G_s}$. This quantity scales the burden in each bin by the typical event length relative to the callable genome, yielding a value proportional to the expected frequency with which events of that class alter tested genomic intervals in that sample.

Finally, we summed these bin-specific rates across bins to obtain sample-level baseline rates $r_{s,t} = \sum_b R_{s,t,b}$ for copy-number increases and decreases. These rates serve as the neutral background rates in the likelihood model and are multiplied by an interval-specific scaled selection coefficient to yield observed CNA fixation across tumors.

#### Selection intensities and segment-level likelihood

For a given sample $s$, the tested intervals may be covered by a single copy-number segment or by multiple segments. At one extreme, one segment may span all tested intervals; at the other, each interval may be covered by a different segment. Samples lacking known copy-number status across all tested intervals were excluded from inference, as were the highly uncommon samples in which a single interval was partially increased in copy-number and partially decreased.

Selection was modeled using interval- and event-type specific selection intensities $\gamma_{i,t} \geq 0$ that multiplicatively scale event rates for segments that overlap selected intervals. The segment-level selection coefficients for increases and decreases are defined as products over the overlapped selected intervals: $g_{s,j,dec} = \prod_{i \in \mathcal{I}(s,j)} \gamma_{i,dec}$ and $g_{s,j,inc} = \prod_{i \in \mathcal{I}(s,j)} \gamma_{i,inc}$, where $\mathcal{I}(s,j)$ is the set of tested intervals overlapped for segment $j$ in sample $s$, and where the product is taken as 1 (selective neutrality) when no interval of a given type overlaps the segment. Given these quantities, the total scaled event rate $v$ for segment $j$ in sample $s$ was defined as $v_{s,j} = g_{s,j,inc} r_{s,inc} + g_{s,j,dec} r_{s,dec}$, where $r_{s,inc}$ and $r_{s,dec}$ are the sample-specific baseline rates for copy-number increases and decreases.

#### Segment-level likelihood under the CNA selection model

Under the model, copy-number events are assumed to arise consistent with a Poisson process such that the probability of a no-copy-change segment is $e^{-v_{s,j}}$. Assuming that non-neutral copy states do not reverse, the probability that at least one event occurs is $1 - e^{-v_{s,j}}$. When at least one event occurs, the observed copy-number state must correspond to a particular event type $t$

and size bin $b$. The probability of each possible event type and size class is proportional to its scaled baseline flux. Specifically, for segment $j$ in sample $s$, the flux associated with event type $t$ and size bin $b$ is $u_{s,j,t,b} = g_{s,j,t} R_{s,t,b}$. Conditional on at least one event occurring on the segment, the probability of observing type $t$ and bin $b$ is $P(t, b|y_{s,j} = 1) = \frac{u_{s,j,t,b}}{v_{s,j}}$. This conditional term defines the segment-level likelihood contribution. For segments with no observed event of any type, the likelihood contribution is $e^{-v_{s,j}}$. For segments with an observed event of type $t$ and bin $b$, the contribution is $(1 - e^{-v_{s,j}})\frac{u_{s,j,t,b}}{v_{s,j}}$, yielding the full likelihood

$$\mathcal{L}_{s,j} = \begin{cases} e^{-v_{s,j}} & \text{if } y_{s,j} = 0, \\ (1 - e^{-v_{s,j}})\ \frac{\mathrm{u}_{s,j,t,b}}{v_{s,j}} & \text{if } y_{s,j} = 1. \end{cases}$$

$$\mathcal{L}(\gamma_{i,t}) = \prod_s \prod_j \mathcal{L}_{s,j}$$

$$\log \mathcal{L}(\gamma_{i,t}) = \sum_s \sum_j \log \mathcal{L}_{s,j}$$

The full log-likelihood is obtained by summing contributions over segments $j$ and samples $s$ ; selection intensities can then be estimated by maximum likelihood using the limited-memory Broyden-Fletcher-Goldfarb-Shanno algorithm.
